# SIMULATING TUMOR MITOCHONDRIAL ENERGETICS THROUGH ENGINEERING-STYLE ENERGY METRICS

**DOI:** 10.1101/2025.09.02.673700

**Authors:** Arturo Tozzi

**Affiliations:** Center for Nonlinear Science, Department of Physics, University of North Texas, Denton, Texas, USA; 1155 Union Circle, #311427 Denton, TX 76203-5017 USA

**Keywords:** oxidative phosphorylation, cristae morphology, tumor microenvironment, immune evasion, metabolic profiling

## Abstract

Cancer progression is linked to alterations in cellular energetics, where malignant cells reprogram their metabolism to sustain proliferation, resist stress and adapt to nutrient limitations. Beyond intrinsic adaptations and plastic dynamic switches between glycolysis and oxidative phosphorylation, recent work has shown that tumors actively remodel their microenvironment by acquiring functional mitochondria from surrounding stromal or immune cells. This process of mitochondrial transfer enhances tumor bioenergetics while simultaneously depleting immune cells of metabolic competence, thereby reinforcing both tumor growth and immune evasion. Despite the role of these mitochondrial exchanges, quantitative frameworks to measure their energetic consequences remain underdeveloped. Conventional assays describe oxygen consumption or glycolytic flux but lack the resolution to capture mitochondrial function in terms of throughput, efficiency and stored energy. To address this limitation, we propose a simulation-based framework that translates engineering-style energy metrics into mitochondrial biology. Specifically, we calculate three parameters that may capture distinct yet complementary aspects of mitochondrial bioenergetics: power density, defined as ATP production per unit mitochondrial volume; surface power density, reflecting ATP production per unit inner membrane area; and energy density, quantifying stored chemical free energy per unit volume. By simulating tumor and immune cell populations before and after mitochondrial transfer, we generate numerical values for each parameter that demonstrate how transfer enhances tumor energetics while diminishing immune function. This provisional quantification establishes a pathway toward standardized bioenergetic biomarkers in cancer, potentially enhancing diagnostic methods and supporting the development of strategies aimed at disrupting pathogenic mitochondrial transfer while restoring metabolic competence to immune cells.

## INTRODUCTION

Cancer energetics is defined by the ability of malignant cells to sustain rapid growth under fluctuating nutrient and oxygen availability, often through reprogramming of metabolic pathways (Loo et al. 2015; Lin et al. 2019; Xia et al. 2021; Pavlova, Zhu, and Thompson 2022; Kubik et al. 2022; Tufail, Jiang, and Li 2024). Central to this process are mitochondria, which not only generate ATP, but also regulate redox balance, biosynthetic precursors and signalling (Ryu et al. 2024). Their adaptive capacity underpins the metabolic plasticity enabling tumor cells to thrive in hostile microenvironments (Yang et al. 2016; Missiroli et al. 2020; Kafkova and Trnka 2020; Guo et al. 2021). More recently, mitochondrial transfer between cells has been identified as a mechanism that not only supports tumor growth by supplying respiration-competent organelles (Borcherding and Brestoff 2023; Watson et al. 2023; Zhang et al. 2023a; Ikeda et al. 2025; Hoover et al. 2025), but also undermines immune surveillance by delivering functionally compromised mitochondria to lymphocytes (Zhang et al. 2023b; Chun, An, and Kim 2025). Current methods used to assess these phenomena rely on indicators like oxygen consumption rates, extracellular acidification or mtDNA content, which, while informative, provide only partial insight into the quantitative performance of mitochondria and their contribution to intercellular dynamics. This underscores the absence of standardized approaches to evaluate mitochondrial activity and the need for comparable measures across diverse cell types and conditions. In response to this limitation, we suggest a simulation that adapts engineering-style energy metrics to mitochondrial biology, with the goal of standardizing quantification of organelle function and transfer effects.

We propose three parameters, informed by physics-based energy metrics, that extend conventional bioenergetic assessments by translating the concepts of power density, surface power density and energy density into the context of mitochondrial biology (Karami-Mosammam et al. 2022; Shang et al. 2023). The three engineering-style energy metrics quantify how energy is produced, transferred or stored relative to physical dimensions such as volume, surface area and time (Kim et al. 2019; Li et al. 2021; Foong et al. 2022). Expressed in standard units like watts per cubic metre or joules per cubic metre, they enable accurate, comparable descriptions of energetic performance across diverse systems:

a. Mitochondrial power density is calculated as the ATP production rate per unit mitochondrial volume, reflecting volumetric throughput.
b. Mitochondrial surface power density is defined as ATP production rate per unit inner membrane or cristae area, thereby linking performance to structural capacity.
c. Mitochondrial energy density captures the total stored chemical free energy per unit mitochondrial volume, integrating ATP, reduced cofactors and the proton motive force.

Together, these parameters establish a multidimensional description of mitochondrial output, efficiency, and reserve that can be applied across diverse experimental systems or simulated environments. In this study, we explicitly calculate their values, providing theoretical benchmarks that anchor these concepts within a quantitative framework for evaluating mitochondrial behaviour in cancer biology

We will proceed as follows. The methodology describes how these metrics are derived and measured within the simulation, then the results section presents their behaviour in the simulated contexts. The subsequent sections develop their interpretation in the setting of tumor–immune interactions. Finally, the discussion evaluates their significance within the broader framework of cancer energetics.

## MATERIALS AND METHODS

We detail a simulation-based approach that integrates mitochondrial geometry, flux kinetics, energy storage and transfer dynamics between tumour and immune cells. Parameters were derived from physical laws and literature-based ranges, with power density, surface power density and energy density computed through explicit mathematical formulations. Our simulation aims to build synthetic mitochondria, assign bioenergetic states and propagate these states through a theoretical tumor–immune co-culture in silico. The simulation time is indexed by discrete steps t = 0,1, …, T with step size Δt. Each cell c contains a set of mitochondria Mc = {m1, …, mNc}. For each mitochondrion m we track volume Vm[m^3^], inner membrane area Am[m^2^], ATP flux n·ATP, m [mol s^−1^], proton motive force components Δψm[V] and ΔpHm and reduced carrier pools nNADH,m[molJ (Imamura et al. 2009; Oncescu and Erickson, 2013; Cox and Smith 2014; Natsubori et al. 2020; Greiner and Glonek 2021). Temperature is fixed at T = 310 K. The ATP free energy is computed as

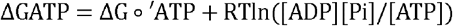

with 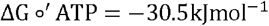. Power per mitochondrion is Pm = n·ATP,m | ΔGATP |. From these primitives we compute:

1. mitochondrial power density pm = Pm/Vm[Wm^−3^],
2. surface power density sm = Pm/Am[Wm^−2^],
3. energy density um = Um/Vm[Jm^−3^], where Um is the stored chemical energy defined below.

Overall, power density describes volumetric throughput of ATP production, surface power density relates ATP synthesis to cristae area and membrane organisation, while energy density captures the static reservoir of chemical free energy (see **Table**). In our simulation, tumor cells are parameterized with higher probabilities of enlarged cristae density, elevated ATP flux capacity, and stronger redox stores, representing their proliferative and anabolic demands documented in cancer metabolism studies. Immune cells are modeled with more moderate volumes, balanced throughput, and greater variability, reflecting their well-established context-dependent energetic adaptability.

**Table.** The three quantitative mitochondrial parameters introduced in our simulation. Each parameter is defined with its physical units and methodological focus, capturing distinct yet complementary dimensions of mitochondrial bioenergetics.

| Parameter | Definition | Methodological focus: what it explains |
| --- | --- | --- |
| **Mitochondrial power density**<br>(W/m<sup>3</sup>) | ATP production rate per unit mitochondrial volume (volumetric throughput of energy conversion) | Explains how much usable power is generated relative to mitochondrial size, capturing overall throughput and enabling comparison of energetic intensity across mitochondria of different volumes. |
| **Mitochondrial surface power density**<br>(W/m<sup>2</sup>) | ATP production rate per unit inner membrane or cristae area (efficiency and packing of oxidative phosphorylation machinery) | Explains how effectively the inner membrane structure supports ATP synthesis, linking functional output to cristae architecture and membrane density. |
| **Mitochondrial energy density**<br>(J/m<sup>3</sup>) | Total stored chemical free energy per unit mitochondrial volume (ATP, NADH, proton motive force) | Explains the reservoir of immediately available energy within a mitochondrion, capturing static storage capacity rather than dynamic throughput. |

### Geometry generator for volume and inner membrane architecture

We need next to formalize mitochondrial geometry and cristae amplification with closed-form area and volume expressions, ensuring that power normalization rests on a reproducible structural basis. Each mitochondrion is represented as a tri-axial ellipsoid with semi-axes (a, b, c) drawn from lognormal priors a ∼ LN(µa, σa^2^) etc., giving volume 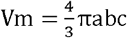. Inner membrane area is generated by a cristae packing model. We define a cristae surface multiplier κ\κ such that Am = κAouter with 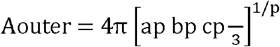 using Knud Thomsen’s approximation with p= 1.6075. The multiplier κ follows a gamma distribution κ ∼ Г(κκ, θκ) to capture tissue variability; typical means are chosen so that Am/Vm = α falls near 10^7^ to 10^8^ in cristae-rich states. Heterogeneity within a cell is produced by drawing I(independently for each mmm conditional on cell type. The geometry generator yields (Vm, Am) for all mitochondria and therefore fixes the structural denominators of pm and sm.

### Flux generator for ATP throughput and redox supply

We specify here the kinetic law connecting membrane area to ATP throughput and embeds temporal variability through a stochastic substrate process, thus establishing a mechanistic source for Pm (Smolders, De Boeck, and Blust 2003; Schmidt, Fisher-Wellman, and Neufer 2021; Bryant and Machta 2023). ATP flux n·ATP, m arises from an Electronic Transport Chain capacity model with Michaelis–Menten saturation and a lognormal random effect (Mok et al. 2024; Singh et al. 2025). Let vmax, m be the maximal ATP synthesis capacity and Sm the effective substrate proxy proportional to oxygen and NADH availability. Flux is

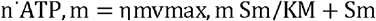

with KM constant across mitochondria and ηm ∼ LN(µη, ση^2^) capturing efficiency dispersion. We set vmax,m = kvAm so that capacity scales with inner membrane area, where kv[mols−^1^m^−2^] is a catalytic surface constant. The substrate proxy evolves as a first order autoregression Sm(t + Δt) = S^−^ + ρ(Sm(t) - S^−^) + σSξt with ξt N(0,1). Power is then Pm = n·ATP, m | ΔGATP | with | ΔGATP | updated from simulated concentrations obeying mass balance d[ATP]/dt = Vm1(n·ATP, m − Jcons, m). Consumption Jcons, m = λm[ATP] uses a linear sink that stabilizes the system.

### Stored energy model for energy density

We provide here explicit thermodynamic formulas for stored energy terms summing to energy density, thereby defining the static counterpart to throughput metrics. Stored energy Um comprises ATP, reduced carriers and the electrochemical proton gradient. We compute UATP,m = nATP,m | ΔGATP | with nATP,m = [ATP]mVmNA and NA Avogadro’s number. Reduced carrier energy uses an equivalent free energy per mole ΔGredox projected onto ATP units to avoid double counting. Let nNADH,m be moles of NADH. Then Uredox,m =nNADH,m∣ΔGredox∣. The proton motive force is Δpm = Δψm - RT/F ln10. ΔpHm and the small-signal membrane energy is UΔψ,m = 21CmAm(Δψm)^2^ with specific capacitance Cm[Fm^−2^]. Buffer energy from ΔpH around equilibrium is approximated by 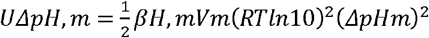 using matrix buffering capacity βH, m[molm^−3^ pH^−1^]. The total energy is Um = UATP, m + Uredox,m+ UΔψ,m+ UΔpH,m and energy density is um = Um/Vm. Concentrations [ATP]m and nNADH,m follow linear Langevin dynamics with reflecting boundaries to maintain physiological ranges.

### Transfer process between cells and state updates

Now transfer needs to be converted into a probabilistic mechanism coupled to energetic states, so that changes in power and energy densities may emerge from exchange dynamics rather than being imposed. Intercellular transfer is simulated as a marked point process on a bipartite graph connecting tumor and immune cells. At each step, a mitochondrion m in donor cell i is selected with probability proportional to a propensity qm = γ0 + γ1pm + γ2um. A recipient j is chosen from the opposite population with probability proportional to wj = *δ*0 + *δ*1p^−^j + *δ*2u^−^j where bars denote cell averages. Transfer occurs with probability π= σ(θ0 + θ1contactij) with σ(x) = (1+e-x)^−1^. On acceptance, *m* is re-indexed to j and its state undergoes adaptation xnew = ϕxold + (1 - ϕ)xenv for x ∈ {Δψ,ΔpH, [ATP], nNADH}, where xenv is the recipient baseline. Contact propensities evolve by a mean-field rule that increases tumor–immune interactions when nutrient fields favor exchange. The nutrient field N is a scalar that diffuses on a lattice with discrete Laplacian Nt + Δt = Nt + DΔ^2^Nt - κCt where Ct is total cellular consumption.

### Definition of biomarkers and aggregation to cell and population scales

We formalize here how single-organelle metrics translate to interpretable cell and population quantities, thus establishing the measurable outputs of the simulation. For each mitochondrion we compute pm, ssm and um at every step. Cell-level biomarkers are arithmetic and area-weighted means: 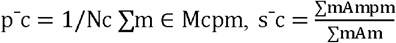 and u^−^c = Nc1∑mum. Population summaries are medians and interquartile ranges of {p ^−^c}\{\bar p_c\}{p^−^c}, {s^−^c}\{\bar s_c\}{s^−^c} and {u^−^c} for tumor and immune groups. We also compute membrane-normalized throughput χc = ∑mPm/∑mAm and volumetric throughput ψc = ∑mPm/∑mVm as cross-checks. Time-aggregated statistics use trapezoidal integration (p^−^) = 1/T ∫ 0Tp^−^c(t)dt approximated discretely. For hypothesis checks we simulate two phases, pre transfer t≤T0 and post transfer t> T0 and compute differences Δp^−^c = p^−^cpost − p^−^cpre, etc.

We calibrate priors so that simulated medians match orders of magnitude reported for mammalian mitochondria: Vm ∼ 10^−18^m3, Am/Vm 10^7^ to 10^8^, n·ATP,m ∼ 10^6^ s^−1^ equivalents (Chapa-Dubocq et al. 2023; Triolo et al. 2023). A synthetic likelihood approach is used. Let z be a vector of target moments and m(Θ) the simulated moments. The synthetic log-likelihood is *ℓ*(Θ) = -21(z - m(Θ))nΣ - 1(z - m(Θ)) with Σ a tuning matrix. Optimization uses L-BFGS over Θ under positivity constraints. Local identifiability is probed by the Fisher Information approximation I(Θ) = JnΣ - 1J, with Jij = *∂*mi/*∂*Θj computed by automatic differentiation on the simulator.

### Statistical comparison and uncertainty quantification

For the simulated cohorts, we compare pre transfer and post transfer phases using Welch statistics with Satterthwaite degrees of freedom. Effect sizes are reported as standardized differences with pooled standard deviation s. Temporal uncertainty is summarized by block bootstrap over contiguous windows to respect autocorrelation. Multiple comparison control uses the Benjamini–Hochberg procedure on p values.

### Numerical implementation and software tools

Simulations are implemented in Python 3.11. Core numerical operations rely on NumPy for array algebra, SciPy for optimization, JAX for automatic differentiation where gradients are required and Numba for just-in-time acceleration of the Monte Carlo transfer kernel. Randomness uses NumPy’s PCG64 bit generator with seed control for reproducibility. Plots are generated using Matplotlib. Units are tracked with Pint to reduce dimensional inconsistency. The solver advances all mitochondria in vectorized form. At each step we update substrate proxies, compute n·ATP,m, calculate power and energy components, then execute transfer proposals using vectorized sampling. Performance is validated by step halving and conservation checks on total carrier pools. The entire workflow is scripted with Snakemake to guarantee deterministic execution from a single configuration file.

Overall, our methods define a coherent workflow to integrate structural modeling, flux generation, energy storage equations and transfer dynamics into a unified simulator.

## RESULTS

We present here the simulation outcomes designed to quantify mitochondrial bioenergetic metrics before and after intercellular transfer. To contextualise the results, we first include an order-of-magnitude worked example for a single mitochondrion, which links geometry, kinetics and thermodynamics to explicit values of power density, surface power density and energy density. This example provides a concrete baseline for interpreting how population-level simulations may reflect systematic changes in tumour and immune cells. Following this reference point, we describe the distributions and statistical comparisons of these metrics across groups, highlighting measurable and significant mitochondrial shifts that emerge within the simulated tumour microenvironment.

### Order-of-magnitude worked example for a single mitochondrion

To ground the simulation in physical reality, we first calculate an order-of-magnitude example for a single mitochondrion using established structural and energetic parameters. This example serves as a reference point against which simulated populations of tumour and immune cells can be compared, ensuring that subsequent results remain anchored to physiologically plausible magnitudes.

A single mitochondrion is instantiated with volume V = 1.0 × 10^−18^ m^3^ and inner membrane area A= 5.0 × 10^−12^ m^2^, consistent with an ellipsoid of micron scale and cristae amplification. ATP flux is set to n·ATP = 1.0 × 10^6^ s^−1^ molecules, which equals 1.66 × 10^−18^mols^−1^. With | ΔGATP |= 50kJmol^−1^, the instantaneous power is P = n·ATP| ΔGATP |= 8.3 × 10^−1^ W. Mitochondrial power density is then 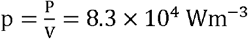 and surface power density is 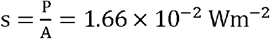. For stored energy we specify [ATP] = 5mM and nNADH = 2.0 × 10^−19^ mol^−19^. The ATP store contributes UATP = [ATP]VNA | ΔGATP | = 2.5 × 10^−13^J. With | ΔGredox |= 220kJmol^−1^, the redox store contributes Uredox = 4.4 × 10^−1^ J. Take Cm = 0.01Fm^−2^ and Δψ = 0.15V, giving UΔψ = 21CmA(Δψ)2= 5.6 × 10^−16^J. With βH = 20molm - 3pH^−1^ and ΔpH = 0.5, the buffer term is 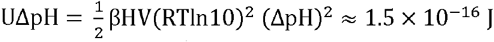. Total stored energy is U ≈ 2.96 × 10^−13^ J and energy density is 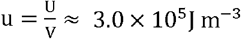.

This grounding not only validates the plausibility of the simulation, but also provides an essential benchmark for interpreting the broader statistical patterns observed in tumour and immune cell populations

### Tumor- and immune-associated mitochondrial metrics

Simulated tumor mitochondria displayed a significant increase in energetic parameters after transfer (Figure). All the parameters studied showed statistical significance with p < 0.001. Mean power density rose from 90,714 W m□^3^ before transfer to 129,699 W m□^3^ after transfer. Surface power density increased from 0.020 W m□^2^ to 0.030 W m□^2^. Energy density increased from 257,217 J m□^3^ to 319,952 J m□^3^. These increments, consistent across replicates, suggest that tumor mitochondria adopt higher volumetric throughput, denser inner membrane efficiency and greater stored energy following acquisition of new organelles.

**Figure.**
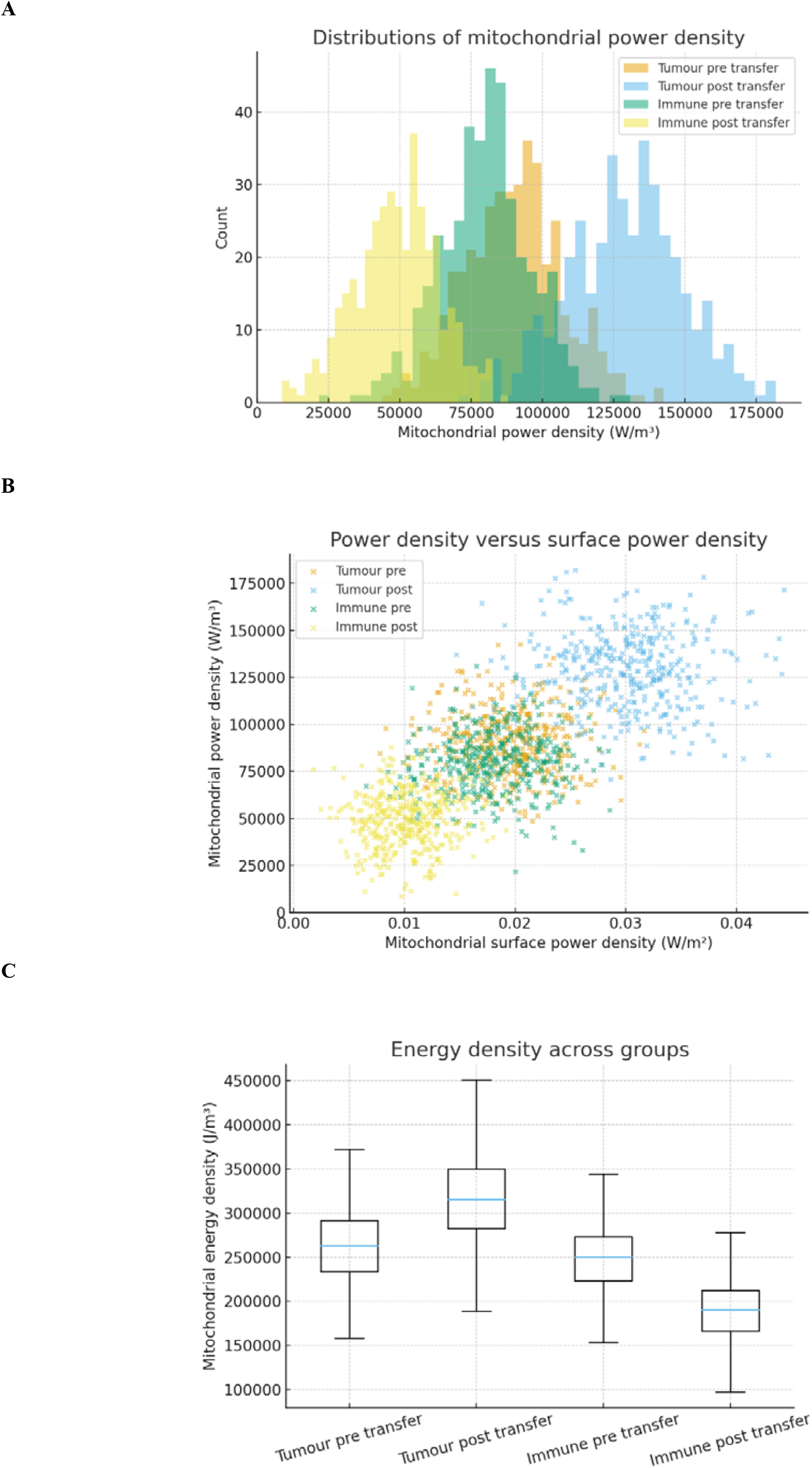
Simulated behaviour of the three mitochondrial bioenergetic metrics, showing their shifts in tumour and immune populations before and after organelle transfer. **Figure A**. Simulated mitochondrial power density distributions in tumor and immune populations before and after organelle transfer. Tumor cells shift toward higher values while immune cells shift lower, consistent with acquisition of high throughput mitochondria by tumors and receipt of impaired organelles by immune cells. Units are W per m^3^. **Figure B**. Power density plotted against surface power density for individual simulated cells. Tumor post transfer points cluster at higher surface power density and higher power density, indicating more efficient and densely packed inner membranes. Immune post transfer points cluster at lower values. Units are W per m^2^ on the x axis and W per m^3^ on the y axis. **Figure C**. Boxplots of mitochondrial energy density across groups. Tumor post transfer shows increased medians relative to tumor pre transfer while immune post transfer shows decreased medians relative to immune pre transfer. Values reflect stored chemical free energy normalized by mitochondrial volume. Units are J per m^3^.

Immune mitochondria displayed a complementary decrease in the same metrics (Figure). Power density dropped from 80,613 W m□^3^ before transfer to 50,112 W m□^3^ after transfer. Surface power density decreased from 0.018 W m□^2^ to 0.010 W m□^2^. Energy density fell from 248,959 J m□^3^ to 192,395 J m□^3^. These consistent declines indicate that immune cell mitochondria acquire reduced capacity, lowered cristae efficiency and diminished energy stores after transfer.

Overall, the juxtaposition of the results observed in tumor and immune cells following simulated intercellular transfer establishes a complementary pattern, in which tumor energetics are enhanced and immune energetics are suppressed, completing the picture of transfer-induced divergence between cell types.

## CONCLUSIONS

By situating mitochondrial energetic changes within the framework of quantitative engineering-style metrics, our simulation revealed quantifiable differences in mitochondrial bioenergetic parameters following intercellular transfer between tumor and immune populations. Tumor cells displayed increases in power density, surface power density and energy density, possibly corresponding to heightened volumetric throughput, enhanced efficiency of inner membrane organization and elevated reservoirs of stored chemical energy. Immune cells, in contrast, exhibited decreases in the same three metrics, possibly indicating diminished bioenergetic capacity, lower cristae efficiency and reduced stored energy after receiving impaired organelles. This outcome supports the feasibility of using these three metrics as standardized indicators of cellular bioenergetic state under conditions mimicking tumor–immune interactions. The meaning of our approach lies in the redefinition of mitochondrial analysis through three interconnected but distinct metrics that quantify throughput, surface efficiency and stored capacity. The three parameters provide a language that translates cellular metabolism into physical parameters with explicit units, making it possible to evaluate intracellular energetic states within a unified framework.

Conventional bioenergetic analysis often relies on oxygen consumption rate measurements, which provide valuable information about mitochondrial activity but capture only one aspect of throughput (Sheeley et al. 2022; Halma, Tuszynski, and Marik 2023; Liu et al. 2024). Similarly, extracellular acidification rates reflect glycolytic contributions but are decoupled from mitochondrial volume or membrane organization (Mookerjee et al. 2017). Fluorescent indicators of membrane potential and pH gradients yield insights into specific components of the proton motive force, but these data are seldom aggregated into a single coherent measure of stored energy (Perry et al. 2011; Weiß and Bohrmann 2019). Electron microscopy supplies detailed structural images of cristae but cannot alone indicate functional throughput. By contrast, our simulation explicitly binds these multiple dimensions into three metrics that can be cross-compared, each with a clear physical unit. Also, the simulation formalizes equations linking substrate flux, membrane architecture and chemical storage, avoiding reliance on isolated descriptors. While other approaches provide partial views, our framework integrates them into a single language that captures throughput, structural efficiency and reserve capacity simultaneously.

Therefore, compared to existing approaches, our framework could discriminate whether observed changes in bioenergetics stem from higher mitochondrial content, improved inner membrane packing or altered energetic reserves, which may be crucial in characterizing tumor metabolism and its impact on immune competence. Still, our simulation demonstrates how engineering principles can be adapted to biological systems, providing a bridge between cellular bioenergetics and physical metrics that are broadly interpretable.

However, limitations must be acknowledged. Being a simulation, all quantitative results are based on parameter distributions drawn from literature estimates, not direct empirical measurements. While the magnitudes and trends are representative, they cannot replace the biological variability observed in living systems. The simplification of mitochondria as ellipsoids with parameterized cristae density, while mathematically tractable, does not capture the full structural complexity of dynamic inner membrane remodeling. ATP flux was modeled using generalized Michaelis– Menten kinetics, which abstracts away the complexity of multi-step enzymatic reactions and regulatory mechanisms. Furthermore, our simulation treated mitochondrial transfer as a probabilistic process governed by energetic propensity, whereas in vivo transfer occurs through multiple physical routes, including tunneling nanotubes and extracellular vesicles, which introduce additional heterogeneity (Guan et al. 2024). The model also assumes uniform environmental conditions, neglecting gradients of oxygen, glucose or pH that strongly influence bioenergetics in tumors. Finally, statistical analyses rely on synthetic populations generated under the assumptions embedded in the model, which limits the generalizability of the significance levels beyond simulation.

Several potential applications, directions for future research and testable experimental hypotheses can be articulated. The metrics themselves suggest new avenues for characterizing mitochondrial performance in cell cultures and patient-derived samples, as oxygen flux, inner membrane architecture and ATP concentration data can be mapped onto the framework. Future research could integrate imaging-based structural quantification with high-resolution respirometry and metabolomics to empirically validate the three parameters in real biological contexts. Testable hypotheses include the prediction that tumors displaying high average mitochondrial power density correlate with aggressive clinical progression or that immune cells exhibiting reduced energy density after mitochondrial transfer are correlated with diminished responsiveness to immunotherapy. Experiments could also assess whether interventions preventing mitochondrial transfer preserve higher immune surface power density, measurable by respirometry combined with ultrastructural imaging. Still, these metrics could be incorporated into standard bioenergetic profiling to assess cell populations in the tumor milieu.

In summary, we described mitochondrial bioenergetics through explicit physical metrics to provide an accurate and reproducible language for assessing energy-related dynamics of tumor microenvironment. The main research question was whether mitochondrial power density, surface power density and energy density can serve as quantitative biomarkers to describe the consequences of intercellular mitochondrial transfer in cancer. While rooted in simulation, our approach suggests the feasibility of standardizing mitochondrial bioenergetics through physically interpretable parameters for the study of cancer metabolism.

## DECLARATIONS

### Ethics approval and consent to participate

This research does not contain any studies with human participants or animals performed by the Author.

### Consent for publication

The Author transfers all copyright ownership, in the event the work is published. The undersigned author warrants that the article is original, does not infringe on any copyright or other proprietary right of any third part, is not under consideration by another journal and has not been previously published.

### Availability of data and materials

All data and materials generated or analyzed during this study are included in the manuscript. The Author had full access to all the data in the study and took responsibility for the integrity of the data and the accuracy of the data analysis.

## Acknowledgements

none.

## Authors’ contributions

The Author performed: study concept and design, acquisition of data, analysis and interpretation of data, drafting of the manuscript, critical revision of the manuscript for important intellectual content, statistical analysis, obtained funding, administrative, technical and material support, study supervision.

## Declaration of generative AI and AI-assisted technologies in the writing process

During the preparation of this work, the author used ChatGPT 4o to assist with data analysis and manuscript drafting and to improve spelling, grammar and general editing. After using this tool, the author reviewed and edited the content as needed, taking full responsibility for the content of the publication.

## REFERENCES

1) Borcherding, Nicholas, and Jordan R. Brestoff. 2023. “The Power and Potential of Mitochondria Transfer.” Nature 623 (7986): 283–91. 10.1038/s41586-023-06537-z Bryant, Samuel J., and Benjamin B. Machta. 2023. “Physical Constraints in Intracellular Signaling: The Cost of Sending a Bit.” Physical Review Letters 131 (6): 068401. 10.1103/PhysRevLett.131.068401.

2) Chapa-Dubocq, Xavier R., Keishla M. Rodríguez-Graciani, Nelson Escobales, and Sabzali Javadov. 2023. “Mitochondrial Volume Regulation and Swelling Mechanisms in Cardiomyocytes.” Antioxidants (Basel) 12 (8): 1517. 10.3390/antiox12081517

3) Chun, Soohyun, Jin An, and Man S. Kim. 2025. “Mitochondrial Transfer between Cancer and T Cells: Implications for Immune Evasion.” Antioxidants 14 (8): 1008. 10.3390/antiox14081008

4) Cox, Brian N., and David W. Smith. 2014. “On Strain and Stress in Living Cells.” Journal of the Mechanics and Physics of Solids 71 (November): 239–52. 10.1016/j.jmps.2014.07.001.

5) Foong, Yee Wei, Javad Shirani, Shuaishuai Yuan, Christopher A. Howard, and Kirk H. Bevan. 2022. “Towards a Pseudocapacitive Battery: Benchmarking the Capabilities of Quantized Capacitance for Energy Storage.” PRX Energy 1 (June): 013007. 10.1103/PRXEnergy.1.013007.

6) Greiner, Jack V., and Thomas Glonek. 2021. “Intracellular ATP Concentration and Implication for Cellular Evolution.” Biology (Basel) 10 (11): 1166. 10.3390/biology10111166.

7) Guan, Fang, Xue Wu, Jiahui Zhou, Yutong Lin, Yaqiong He, Chao Fan, Zhihui Zeng, and Wenwen Xiong. 2024. “Mitochondrial Transfer in Tunneling Nanotubes—A New Target for Cancer Therapy.” Journal of Experimental & Clinical Cancer Research 43 (1): 147. 10.1186/s13046-024-03069-w.

8) Guo, Xiaojun, Ning Yang, Weiqiang Ji, Hao Zhang, Xiaoli Dong, Zheng Zhou, Lei Li, Han-Ming Shen, Shao Q. Yao, and Weiping Huang. 2021. “Mito-Bomb: Targeting Mitochondria for Cancer Therapy.” Advanced Materials 33 (43): e2007778. 10.1002/adma.202007778.

9) Hoover, Garrett, Sophie Gilbert, Owen Curley, Claire Obellianne, Michael T. Lin, William Hixson, Thomas W. Pierce, et al. 2025. “Nerve-to-Cancer Transfer of Mitochondria during Cancer Metastasis.” Nature 644 (8075): 252–62. 10.1038/s41586-025-09176-8.

10) Ikeda, Hiroshi, Koji Kawase, Takashi Nishi, Yuka Tanaka, Hideki Yamamoto, Shinya Matsuda, Rie Nakamura, et al. 2025. “Immune Evasion through Mitochondrial Transfer in the Tumour Microenvironment.” Nature 638 (8034): 225–36. 10.1038/s41586-024-08439-0.

11) Imamura, Hiromi, Kim P. Huynh Nhat, Hiroko Togawa, Akira Saito, Ryota Iino, Yuichi Kato-Yamada, Takeharu Nagai, and Hiroyuki Noji. 2009. “Visualization of ATP Levels inside Single Living Cells with Fluorescence Resonance Energy Transfer-Based Genetically Encoded Indicators.” Proceedings of the National Academy of Sciences 106 (37): 15651–56. 10.1073/pnas.0904764106.

12) Kafkova, Anna, and Jiri Trnka. 2020. “Mitochondria-Targeted Compounds in the Treatment of Cancer.” Neoplasma 67 (3): 450–60. 10.4149/neo_2020_190725N671.

13) Karami-Mosammam, Morteza, Dominik Danninger, Dominik Schiller, and Martin Kaltenbrunner. 2022. “Stretchable and Biodegradable Batteries with High Energy and Power Density.” Advanced Materials 34 (32): e2204457. 10.1002/adma.202204457.

14) Kim, Minjeong, Dongwoo Park, Md. Mahbub Alam, Seungwan Lee, Pilju Park, and Jaehoon Nah. 2019. “Remarkable Output Power Density Enhancement of Triboelectric Nanogenerators via Polarized Ferroelectric Polymers and Bulk MoS□ Composites.” ACS Nano 13 (4): 4640–46. 10.1021/acsnano.9b00750.

15) Kubik, Joanna, Ewa Humeniuk, Grzegorz Adamczuk, Beata Madej-Czerwonka, and Anna Korga-Plewko. 2022. “Targeting Energy Metabolism in Cancer Treatment.” International Journal of Molecular Sciences 23 (10): 5572. 10.3390/ijms23105572.

16) Li, Zhen, Chao Xin, Yiqing Peng, Ming Wang, Jiajie Luo, Shuang Xie, and Hong Pu. 2021. “Power Density Improvement of Piezoelectric Energy Harvesters via a Novel Hybridization Scheme with Electromagnetic Transduction.” Micromachines (Basel) 12 (7): 803. 10.3390/mi12070803.

17) Lin, Chun, Thomas C. Salzillo, Daniel A. Bader, Samuel R. Wilkenfeld, Dana Awad, Tiffany L. Pulliam, Pradip Dutta, et al. 2019. “Prostate Cancer Energetics and Biosynthesis.” In Advances in Experimental Medicine and Biology, vol. 1210, 185–237. 10.1007/978-3-030-32656-2_10.

18) Loo, Jamie M., Amy Scherl, Anh Nguyen, Foon Yee Man, Eric Weinberg, Zhihui Zeng, Leonard Saltz, Philip B. Paty, and Sohail F. Tavazoie. 2015. “Extracellular Metabolic Energetics Can Promote Cancer Progression.” Cell 160 (3): 393–406. 10.1016/j.cell.2014.12.018.

19) Missiroli, Sabrina, Marco Perrone, Isabella Genovese, Paolo Pinton, and Carlotta Giorgi. 2020. “Cancer Metabolism and Mitochondria: Finding Novel Mechanisms to Fight Tumours.” EBioMedicine 59 (September): 102943. 10.1016/j.ebiom.2020.102943.

20) Mok, Darren Z. L., Danny J. H. Tng, Jia Xin Yee, Valerie S. Y. Chew, Christine Y. L. Tham, Justin S. G. Ooi, Hwee Cheng Tan, et al. 2024. “Electron Transport Chain Capacity Expands Yellow Fever Vaccine Immunogenicity.” EMBO Molecular Medicine 16 (6): 1310–23. 10.1038/s44321-024-00065-7

21) Mookerjee, Shona A., Ákos A. Gerencser, David G. Nicholls, and Martin D. Brand. 2017. “Quantifying Intracellular Rates of Glycolytic and Oxidative ATP Production and Consumption Using Extracellular Flux Measurements.” Journal of Biological Chemistry 292 (17): 7189–7207. 10.1074/jbc.M116.774471.

22) Natsubori, Akinori, Tetsuya Tsunematsu, Akiko Karashima, Kenji F. Tanaka, Akihiro Takahashi, Yuki Takata, and Hiroki Ueda. 2020. “Intracellular ATP Levels in Mouse Cortical Excitatory Neurons Vary with Sleep– Wake States.” Communications Biology 3 (1): 491. 10.1038/s42003-020-01215-6.

23) Xia, Li, Ling Oyang, Jian Lin, Song Tan, Yajuan Han, Ning Wu, Ping Yi, et al. 2021. “The Cancer Metabolic Reprogramming and Immune Response.” Molecular Cancer 20 (1): 28. 10.1186/s12943-021-01316-8.

24) Yang, Yumei, Svetlana Karakhanova, Wolfram Hartwig, Joerg G. D’Haese, Petr P. Philippov, Jens Werner, and Alexander V. Bazhin. 2016. “Mitochondria and Mitochondrial ROS in Cancer: Novel Targets for Anticancer Therapy.” Journal of Cellular Physiology 231 (12): 2570–81. 10.1002/jcp.25349.

25) Oncescu, Vlad, and David Erickson. 2013. “High Volumetric Power Density, Non-Enzymatic, Glucose Fuel Cells.” Scientific Reports 3 (March): 1226. 10.1038/srep01226.

26) Pavlova, Natalya N., Jun Zhu, and Craig B. Thompson. 2022. “The Hallmarks of Cancer Metabolism: Still Emerging.” Cell Metabolism 34 (3): 355–77. 10.1016/j.cmet.2022.01.007.

27) Perry, Seth W., John P. Norman, Justin Barbieri, Edward B. Brown, and Harris A. Gelbard. 2011. “Mitochondrial Membrane Potential Probes and the Proton Gradient: A Practical Usage Guide.” BioTechniques 50 (2): 98–115. 10.2144/000113610

28) Ryu, Keun Woo, Tak Shun Fung, Daphne C. Baker, Michelle Saoi, Jinsung Park, Christopher A. Febres-Aldana, Rania G. Aly, et al. 2024. “Cellular ATP Demand Creates Metabolically Distinct Subpopulations of Mitochondria.” Nature 635 (8039): 746–54. 10.1038/s41586-024-08146-w.

29) Schmidt, Cameron A., Kelsey H. Fisher-Wellman, and P. Darrell Neufer. 2021. “From OCR and ECAR to Energy: Perspectives on the Design and Interpretation of Bioenergetics Studies.” Journal of Biological Chemistry 297 (4): 101140. 10.1016/j.jbc.2021.101140.

30) Shang, Yuchen, Chao Li, Guo Yu, Yong Yang, Wenhao Zhao, and Weiguo Tang. 2023. “High Storable Power Density of Triboelectric Nanogenerator within Centimeter Size.” Materials (Basel) 16 (13): 4669. 10.3390/ma16134669.

31) Singh, Deepak, Thomas Robin, Michael Urbakh, and colleagues. 2025. “High-Order Michaelis-Menten Equations Allow Inference of Hidden Kinetic Parameters in Enzyme Catalysis.” Nature Communications 16: 2739. 10.1038/s41467-025-57327-2

32) Smolders, Roel, Gudrun De Boeck, and Ronny Blust. 2003. “Changes in Cellular Energy Budget as a Measure of Whole Effluent Toxicity in Zebrafish (Danio rerio).” Environmental Toxicology and Chemistry 22 (4): 890–99. 10.1897/1551-5028(2003)022<0890:CCEBAM>2.0.CO;2.

33) Triolo, Matthew, Steven Wade, Nicole Baker, and Mireille Khacho. 2023. “Evaluating Mitochondrial Length, Volume, and Cristae Ultrastructure in Rare Mouse Adult Stem Cell Populations.” iScience Protocols (February 17): 102107. 10.1016/j.xpro.2023.102107

34) Tufail, Muhammad, Chun Hua Jiang, and Nan Li. 2024. “Altered Metabolism in Cancer: Insights into Energy Pathways and Therapeutic Targets.” Molecular Cancer 23 (1): 203. 10.1186/s12943-024-02119-3.

35) Watson, Douglas C., Didem Bayik, Sigrid Storevik, Sebastian S. Moreino, Samuel A. Sprowls, Jie Han, Maria T. Augustsson, et al. 2023. “GAP43-Dependent Mitochondria Transfer from Astrocytes Enhances Glioblastoma Tumorigenicity.” Nature Cancer 4 (5): 648–64. 10.1038/s43018-023-00556-5.

36) Weiß, Ina, and Johannes Bohrmann. 2019. “Electrochemical Patterns during Drosophila Oogenesis: Ion-Transport Mechanisms Generate Stage-Specific Gradients of pH and Membrane Potential in the Follicle-Cell Epithelium.” BMC Developmental Biology 19 (1): 12. 10.1186/s12861-019-0192-x

37) Zhang, Hong, Xiaoqing Yu, Jinxia Ye, Hao Li, Jing Hu, Yanyan Tan, Ying Fang, et al. 2023a. “Systematic Investigation of Mitochondrial Transfer between Cancer Cells and T Cells at Single-Cell Resolution.” Cancer Cell 41 (10): 1788–802.e10. 10.1016/j.ccell.2023.09.003.

38) Zhang, Hongyi, Xuexin Yu, Jianfeng Ye, Huiyu Li, Jing Hu, Yuhao Tan, Yan Fang, Esra Akbay, Fulong Yu, Chen Weng, Vijay G. Sankaran, Robert M. Bachoo, Elizabeth Maher, John Minna, Anli Zhang, and Bo Li. 2023b. “Systematic Investigation of Mitochondrial Transfer between Cancer Cells and T Cells at Single-Cell Resolution.” Cancer Cell 41 (10): 1788–802.e10. 10.1016/j.ccell.2023.09.003

